# Socio-economic disadvantage is associated with alterations in brain wiring economy

**DOI:** 10.1101/2022.06.08.495247

**Authors:** Roma Siugzdaite, Danyal Akarca, Amy Johnson, Sofia Carozza, Alexander L Anwyl-Irvine, Stepheni Uh, Tess Smith, Giacomo Bignardi, Edwin Dalmaijer, Duncan E. Astle

**Author notes:** Corresponding author: (RS). Authors contributed equally to this work (Roma Siugzdaite, Danyal Akarca, Duncan E. Astle).

## Abstract

The quality of a child’s social and physical environment is a key influence on brain development, educational attainment and mental wellbeing. However, there still remains a mechanistic gap in our understanding of how environmental influences converge on changes in the brain’s developmental trajectory. In a sample of 145 children with structural diffusion tensor imaging data, we used generative network modelling to simulate the emergence of whole brain network organisation. We then applied data-driven clustering to stratify the sample according to socio-economic disadvantage, with one of the resulting clusters containing mostly children living below the poverty line. A formal comparison of the simulated networks from the generative model revealed that the computational principles governing network formation were subtly different for children experiencing socio-economic disadvantage, and that this resulted in significantly altered developmental timing of network modularity emergence. Children in the low socio-economic status (SES) group had a significantly slower time to peak modularity, relative to the higher SES group (*t*_(69)_ = 3.02, *P* = 3.50 × 10^-4^, d = 0.491). In a subsequent simulation we showed that the alteration in generative properties increases the variability in wiring probabilities during network formation (KS test: D = 0.012, *P* < 0.001). One possibility is that multiple environmental influences such as stress, diet and environmental stimulation impact both the systematic coordination of neuronal activity and biological resource constraints, converging on a shift in the economic conditions under which networks form. Alternatively, it is possible that this stochasticity reflects an adaptive mechanism that creates “resilient” networks better suited to unpredictable environments.

**Author Summary:** We used generative network models to simulate macroscopic brain network development in a sample of 145 children. Within these models, network connections form probabilistically depending on the estimated “cost” of forming a connection, versus topological “value” that the connection would confer. Tracking the formation of the network across the simulation, we could establish the changes in global brain organisation measures such as integration and segregation. Simulations for children experiencing socio-economic disadvantage were associated with a shift in emergence of a topologically valuable network property, namely modularity.

## Introduction

Over the first three decades of life the human brain undergoes a gradual process of macroscopic organisation [1, 2], with the emergence and refinement of large-scale networks [3, 4].These networks have a number of important characteristics, like heavy-tailed distributions and densely connected hub regions [5, 6]. This organisation can be captured at the level of an individual as a structural ‘connectome’ – a comprehensive model of whole-brain connectivity for that person. The emergence of connectome organisation over development is non-linear, varies across individuals, and has been linked to multiple aspects of cognition and behaviour. For example, variability in the integration of nodes within the frontal lobes is significantly associated with executive function difficulties in childhood and adolescence [7], and difficulties in cognition during childhood have been linked to significant differences in particular topological features of the connectome [8].

Connectome organization reflects the outcome of an economic trade-off between minimising wiring costs and allowing the emergence of functionally valuable patterns of connectivity between multiple neuronal populations [9]. In short, the organisation of a connectome likely reflects the maximisation of a particular network topology, within the context of resource constraints. The energetic and metabolic cost of each connection can be approximated by the anatomical distance it spans – long connections may enhance the overall organisation of the connectome [10], but they are costly to form and maintain. Across development, as new connections are made, this trade-off between ‘cost’ and topological ‘value’ is continually negotiated [11]. This can be modelled computationally using Generative Network Modelling (GNM), a biophysically embedded computational model that simulates the development of large-scale neural networks [12, 13]. The first instantiation of these models focussed on cost minimisation [14] and subsequently incorporated topological value [12]. The addition of an energy equation, explicitly comparing simulations to observed networks on key global topological distributions, allowed for these models to be optimised [13]. The model applied in the current paper formalises the trade-off between costs and topological values, and, by calibrating the relative balance between these parameters, produces a time-resolved probabilistic model of network formation [12, 13]. This computational approach is in its infancy, but so far there is one consistent result: if the topological ‘value’ term in the model reflects the homophily principle, then we can successfully simulate realistic connectomes. Tiny adjustments to the trade-off between ‘cost’ and ‘value’ are needed to capture each person’s connectome, implying that the wiring economy is slightly different for each person, and that this provides a parsimonious account of the individual variability observed in connectome organisation [11].

Homophily is simply defined as a preference for similarity. In the context of networks, similarity is typically computed in terms of overlapping network topology, such as common neighbours or overlapping connection patterns, such that regions which overlap in their respective connectivity are more likely to wire together. GNMs that incorporate this in the economic trade-off routinely produce the best simulations of structural and functional connectomes [15, 12]. One interpretation is that homophily is simply a macroscopic manifestation of Hebbian learning – “those that fire together, wire together” [12, 16] – and therefore reflects microstructural connectivity played out on a macroscopic scale. When networks are grown according to this principle, within a distance-penalising biophysical framework, the simulation is a close match to a real connectome.

But how does the environment in which development takes place shape this economic trade-off? Multiple environmental factors have a causal influence on cognitive and behavioural outcomes [17 –21], with long-term consequences for mental and physical health [22 – 24]. This influence is sometimes captured by the metric ‘socio-economic status’ (SES). SES is calculated in various ways, typically incorporating parental education level, occupation and household income [25, 26], but likely reflects a much wider set of more proximal environmental influences, like diet, housing conditions, school quality, parental mental health, or access to outside space [18, 27 – 30].

Socio-economic disadvantage is also associated with brain development [17, 18, 31, 32, 33]. Many studies have focused on the structural and functional integrity of particular brain structures, such as the hippocampus [34–37], amygdala [34, 35, 38, 36] and prefrontal cortex [36]. But global brain organisation also covaries with SES [39–43]. An emerging popular theoretical account is that socio-economic disadvantage alters the trajectory of whole-brain development, possibly via a stress responsivity mechanism [42]. Accordingly, negative and repeated experiences may accelerate brain development and reduce plasticity, leading to early maturation of the brain networks. Early disadvantage may curtail the cycle of synaptic proliferation, myelination and pruning that is thought to gradually enhance network organisation [44]. In the short term, these changes can result in what looks like accelerated brain development, with more rapid cortical thinning. However, this shortened trajectory may result in less optimal network organisation in mid-late childhood and adolescence. By contrast, a slower developmental trajectory may prolong the maturation process [42] and maximise the emergence of valuable topological characteristics within the network [45].

One possibility is that socio-economic disadvantage is significantly associated with a systematic shift in neural wiring economy, thereby adjusting the relative trade-off between the cost of forming a connection and topological value it adds to the network. We test this possibility in the current study. We analysed data from 161 children aged 6.8 to 12.8 years with targeted recruitment to lower SES households, such that 33% of children were growing up below the official poverty line in the UK. Diffusion tensor imaging (DTI) data were used create individual connectomes for each child. Individual GNMs then simulated the growth of each child’s connectome, such that we could formally compare wiring rules and optimise parameters for each child. These parameters then enabled us to explore the trajectories of simulated network formation over time, relate them to other properties of the developing brain, and crucially test how the economics of network formation is associated with childhood SES.

## Results

### Simulating the structural connectomes

Individual connectomes for 161 children were created using DTI structural scans (details of connectome construction can be found in the Methods, *Brain network creation and measurements* section). We parcellated structural images according to Brainnetome atlas with 246 ROIs and quantified connectivity between ROIs using streamline tractography. We then modelled each connectome using Generative Network Modelling (GNM) for a sample of 161 participants, 145 of these participants had near-complete SES data and were thus taken into the subsequent steps of the analysis.

At the heart of a GNM is a simple wiring equation [12, 13]. Across all 246 ROIs, at each iteration, the wiring equation calculates which two regions will connect. This calculation is determined by trading-off the cost of a connection forming, against the potential value of the connection being formed. This produces a probability matrix, which updates as the simulation proceeds, at each iteration determining the next connection to form. The equation can be expressed as:

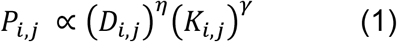

where *D_i,j_* represents the Euclidean distance between nodes *i* and *j* (i.e., “costs”), and *K_i,j_* reflects the value (i.e., “attractiveness”) in forming a connection. *P_i,j_* represents the wiring probability as a function of the product of the parameterized costs and values. *D_i,j_* is parameterized by the scalar *η*, which changes how distances influence the probability of a connection forming. This is traded-off against *K_i,j_*, which represents some relationship between nodes, which can be thought of as a topological value (or “rule”) driving node *i* to connect with node *j. K_i,j_* is parameterized by a distinct scalar *γ*. The parameter combinations for η and γ are selected using a Voronoi tessellation procedure (described in the Methods, *Model fitting with the Voronoi tessellation procedure* section). *K_i,j_* can take a range of different forms and can, in principle, be selected from any non-geometric growth rule used to model social and economic networks. In our analysis we fit 13 different terms for K (the details for these 13 rules can be found in the Methods, *Generative network model to grow connectomes* section), each time trading the respective topological property against the biophysical cost penalty.

The GNM simulates connectome formation for each child in our sample, using the 13 different rules, by adding additional connections until the overall number of connections in the simulation matches those in the respective observed connectome. To test the quality of the simulation an energy function, *E*, calculates the dissimilarity between simulated and observed networks [13].

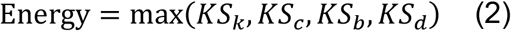

where KS is the Kolmogorov–Smirnov statistic comparing degree *k*, clustering coefficient *c*, betweenness centrality *b*, and edge length *e* distributions of simulated and observed networks. Minimizing *E* finds parameters *η* and *γ* which generate networks most closely approximating the observed network. These four well-known measures that have been used within simulated graph modelling [46, 47, 48].

In summary, the GNM simulates the formation of networks for each child in our sample, according to a different trade-off, using 13 different generative rules. Different economic contexts are produced using the *η* and *γ* scalars to calibrate different relative trade-offs between the distance penalty and value term in the wiring equation. The quality of the simulation is then determined by calculating its energy [13].

### Homophily-based generative rules best simulate network emergence

We ranked each simulation according to its fit, relative to each participant’s observed streamline connectome. Across the sample, connectomes simulated according to homophily rules achieve significantly less energy than all the non-homophily rules (for example, the values ‘matching’ vs ‘c-average’ *t*_(320)_ = −66.01, *P* < 0.001, vs ‘c-minimum’ *t*_(320)_ = 83.9, *P* < 0.001, vs ‘c-max’) *t*_(320)_ = −64.60, *P* < 0.001, vs ‘c-diff’ *t*_(320)_ = −77.47, *P* < 0.001, vs ‘c-product’ *t*_(320)_ = −84.86, *P* < 0.001, vs ‘degree-ave’ *t*_(320)_ = −63.38, *P* < 0.001, vs ‘degree-min’ *t*_(320)_ = − 73.6, P < 0.001, vs ‘degree-max’ *t*_(320)_ = −59.41, *P* < 0.001, vs ‘degree-diff’ *t*_(320)_ = −56.66, *P* < 0.001, vs ‘spatial’ *t*_(320)_ = −85.01, *P* < 0.001) (Fig 1f, Table S2). The best performing of the 13 rules is ‘matching’, which favours connections between nodes with a high normalized overlap in neighbourhoods. Put simply, if nodes ‘A’ and ‘B’ have similar neighbourhoods, then a connection between them is seen as highly valuable, and is likely to form despite the distance penalty.

**Fig 1.**
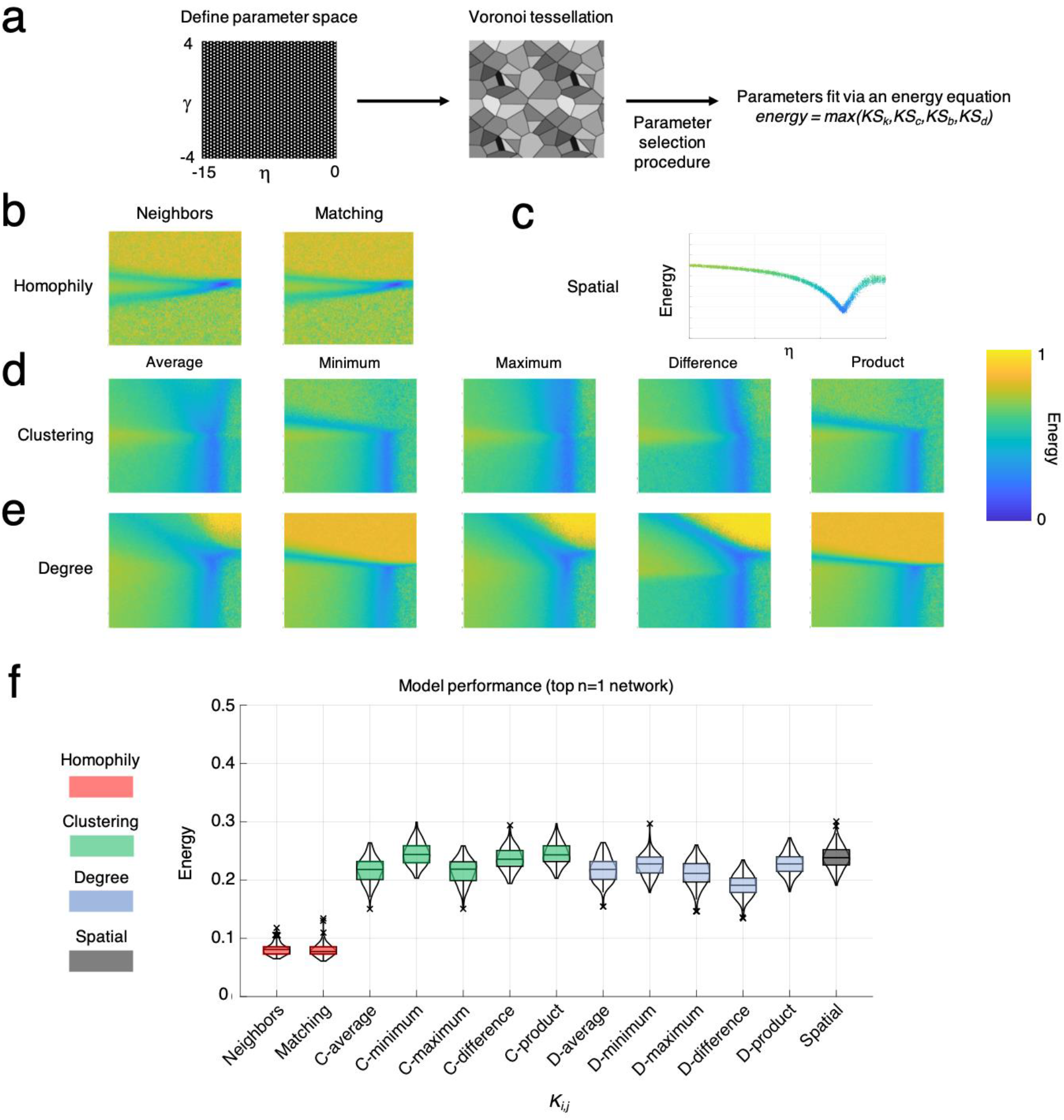
Generative network modelling across thirteen wiring rules shows the homophily produces simulations most similar to those observed. (**a**) We defined the limits of the parameter space as η [-15, 0] and γ [-4, 4], before running a Voronoi tessellation procedure which samples the space in iterations proportionally to the model fit, defined by the energy equation. (**b**) The energy landscapes of homophily-based rules, neighbors and matching. These models simulate network development according to overlaps in connectivity structure. Each landscape plot includes all n=10,000 simulations from all n=161 subjects (n=1,600,000 simulations) overlaid on each other. (**c**) The energy of the spatial model across η, which simulates network development only according to the *D_i,j_* term. (**d**) The energy landscapes of clustering-based rules. These models simulate network development according to relationships in terms of clustering coefficients. Each landscape plot includes all n=10,000 simulations from all n=160 subjects (n=1,600,000 simulations) overlaid on each other. (**e**) The energy landscapes of clustering-based rules. These models simulate network development according to relationships in terms of degree. Each landscape plot includes all n=10,000 simulations from all n=160 subjects (n=1,600,000 simulations) overlaid on each other. (**f**) The energy of the top n=1 performing simulated network for each of the n=161 subjects, across thirteen wiring rules used as the *K_i,j_* term. The boxplot presents the median and IQR.

The parameter landscape for each rule depicts the parameter combination that provides the lowest energy (Fig 1b,d,e). Homophily-based rules have the narrowest low energy portion of that landscape, but provide the lowest energy relative to other rules. This implies that homophily-based rules provide the best approximation of a signal that drives real network formation, but that the economic trade-off between this and the cost parameter is tightly calibrated.

Next we focused specifically on the best performing GNM, the “matching” generative model. We performed the same procedure as before but for a greater number of simulations to improve parameter estimation (50,000 simulations in the parameter space −4 ≤ η ≤ 0, 0.2 ≤ γ ≤ 0.5) (S3 Fig.). Due to inherent stochasticity in the generative network model (as connections form probabilistically), for each subject we calculated their averaged top n=1, n=10, n=50, n=100 and n=500 networks with the lowest energy using η (“costs”) and γ (“values”) parameters which produced these networks (S3 Table). In the further analysis we took forward the n=500 high performing wiring combinations of η and γ values. For each network measure, we plotted mean cumulative density functions across all observed versus simulated nodes within each network (Fig 2). Across multiple network measures the distributional properties of the simulated networks track closely those of the observed streamline connectomes. In S2 Fig. these parameters are compared to each other, and to global, nodal and morphological measures of brain structure.

**Fig 2.**
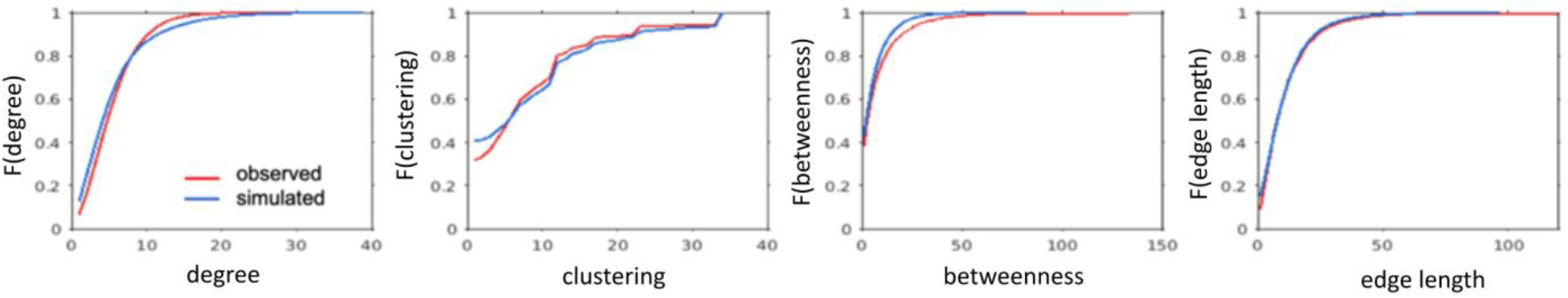
Topological distribution properties for simulated and observed networks based on four topological measures: degree, clustering coefficient, betweenness centrality and edge length.

### Time course of simulated network trajectories

To understand the emergence of global organization during network formation, we identified trajectories of brain segregation and integration during the simulation process. At each moment in the formation of the network we calculated network modularity (via the Q statistic, a measure of segregation) and integration (as inferred by global efficiency). This can be seen in Fig 3. Integration showed a linear change in time, though modularity reached its peak early in network formation before linearly decreasing. For each participant we calculated the time-to-peak and the peak height value of network modularity. Interestingly these two measures are positively correlated (*r* = 0.24, *P* = 4.3×10^-4^; and *r* = 0.253, *P* = 2.70×10^-3^, after the mean framewise displacement is partialled out).

**Fig 3.**
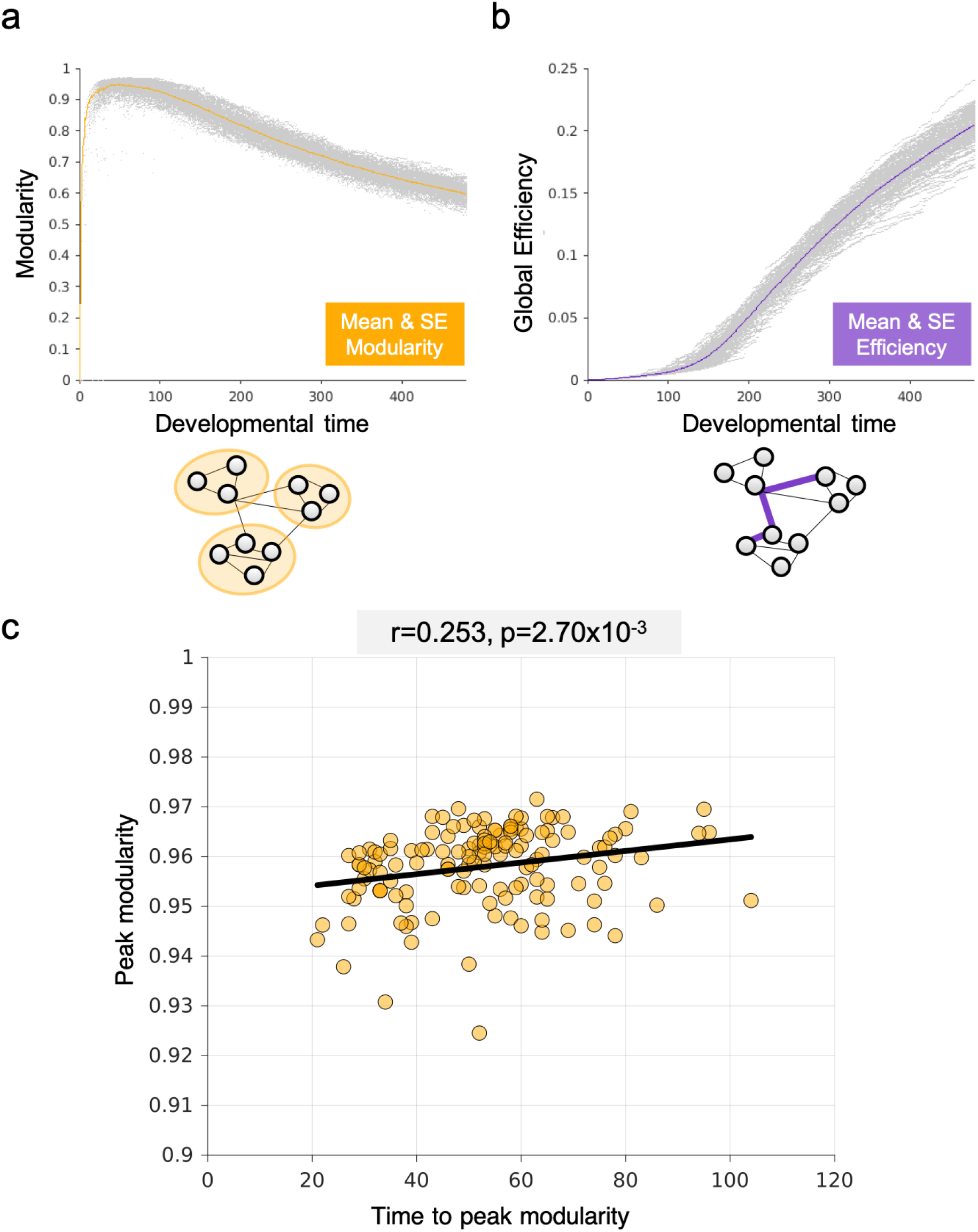
The time course of simulated network statistics over the course of network development. (**a**) Modularity of across all n=140 subjects with SES data, including their mean and SE (yellow line). Light gray points relate to individual points from each subject. Below is a schematic highlighting how modular partitions relate to the segregation of the network. (**b**) Integration of across all n=140 subjects with SES data, including their mean and SE (purple line). Light gray points relate to individual points from each subject. Below is a schematic highlighting how global efficiency captures shortest paths between nodes within the network. (**c**) Taking the peak modularity and time-to-peak modularity, we find a positive correlation (r=0.253, p=2.70×10^-3^; with mean framewise displacement partialled out) suggesting that the later the peak occurs in the generative process, the higher the peak modularity.

### Environmental associations with brain wiring

To assess each child’s socio-economic background, data from six classically used measures of SES were z-scored and entered into a hierarchical clustering algorithm. These included free school meal status [49, 50], equivalised income (income adjusted for household size, [51, 52], neighbourhood deprivation (the Index of Multiple Deprivation, [53] Lang et al 2008), parental occupation status [54], maternal education level [55] and average parental education level [56]. By hierarchically clustering these measures, we obtained a comprehensive index of each child’s SES.

A Silhouette analysis revealed that two clusters provided the best separation within the dataset (Silhouette Coefficient = 0.62). These two clusters are described in Fig 4. The first cluster contained 75 participants, and the second contained 70 participants. We subsampled from the first group in order to produce two identically sized clusters of 70 participants with equivalent average ages (Cluster 1: 9.24±1.22 SD, Cluster 2: 9.58±1.53 SD) and gender split (Cluster 1: 37 girls, 33 boys, Cluster 2: 39 girls, 31 boys). The first cluster broadly captured children from high SES homes, and the second cluster captured children from low SES homes. For instance, in the first group only 1 child was in receipt of free school meals, and under 6% were living below the official poverty line; in the second group 23 children were in receipt of free school meals, and over 60% were living below the official poverty line. Unsurprisingly, these groups differed significantly on the measures included in the original clustering algorithm: children in Cluster 2 were more likely to be in receipt of free school meals (Cluster 1: 1.4%; Cluster 2: 32.9%, *t*_(77.77)_ = −5.4 *P* = 7.3e-07), have lower household income (Cluster 1: 0.48; Cluster 2: 0.2, *t*_(101.36)_ = 9.32, *P* = 2.7e-15), live in more deprived neighbourhoods (Cluster 1: 0.62; Cluster 2: 0.38, *t*_(137.19)_ = 5.98, *P* = 1.8e-08), and have parents with lower occupation status (Cluster 1: 0.67; Cluster 2: 0.38, *t*_(135.87)_ = 4.78, *P* = 4.57e-06), maternal (Cluster 1: 0.81; Cluster 2: 0.31, *t*_(136.2)_ = 15.62, *P* < 2.2e-16) and average parental education level (Cluster 1: 0.73; Cluster 2: 0.37, *t*_(2130.83)_ = 15.2, *P* < 2.2e-16). Cluster membership also generalised to a number of measures that were not included in the original clustering algorithm. These included non-standard measures of SES: Cluster 2 members were more likely to eat more unhealthy foods each day (Cluster 1: 0.14; Cluster 2: 0.27, *t*_(134.54)_ = −5.53, *P* = 1.63e-07), to have poorer health through the life (Cluster 1: 0.89; Cluster 2: 0.78, *t*_(119.45)_ = 3.37, *P* = 0.00098), have a computer or television in their bedroom (Cluster 1: 0.57; Cluster 2: 0.84, *t*_(126.74)_ = −3.67, *P* = 0.0004), spent less time reading (Cluster 1: 0.76; Cluster 2: 0.6, *t*_(138)_ = 3.02, *P* = 0.003), and significantly less likely to encounter other languages in the home (Cluster 1: 0.27; Cluster 2: 0.05, *t*_(104)_ = 3.55, *P* = 0.0006), but no difference in days spent outside (Cluster 1: 0.57; Cluster 2: 0.67, *t*_(136.48)_ = −1.82, *P* = 0.07), and if child has mobile phone or not (Cluster 1: 0.117; Cluster 2: 0.114, *t*(99.72) = 0.1046, *P* = 0.9169). There were also a number of significant differences between the clusters on non-SES related measures. Children in Cluster 2 had significantly higher behavioural difficulties (Cluster 1: 0.23; Cluster 2: 0.34, *t*_(131.88)_ = −3.62, *P* = 0.0004), significantly lower fluid reasoning skills (Cluster 1: 0.58; Cluster 2: 0.39, *t*_(137.97)_ = 5.46, *P* = 2.16e-07), vocabulary (Cluster 1: 0.7; Cluster 2: 0.6, *t*_(136.53)_ = 3.4, *P* = 0.0009) and significantly poorer reading skills (Cluster 1: 0.47; Cluster 2: 0.39, *t*_(137.49)_ = 2.78, *P* = 0.006), though no significant differences in maths performance (Cluster 1: 0.39; Cluster 2: 0.38, *t*_(137.02)_ = 0.46, *P* = 0.6458). With these two well-defined clusters, we next examined the mean parameter combinations needed to best simulate their connectomes.

**Fig 4.**
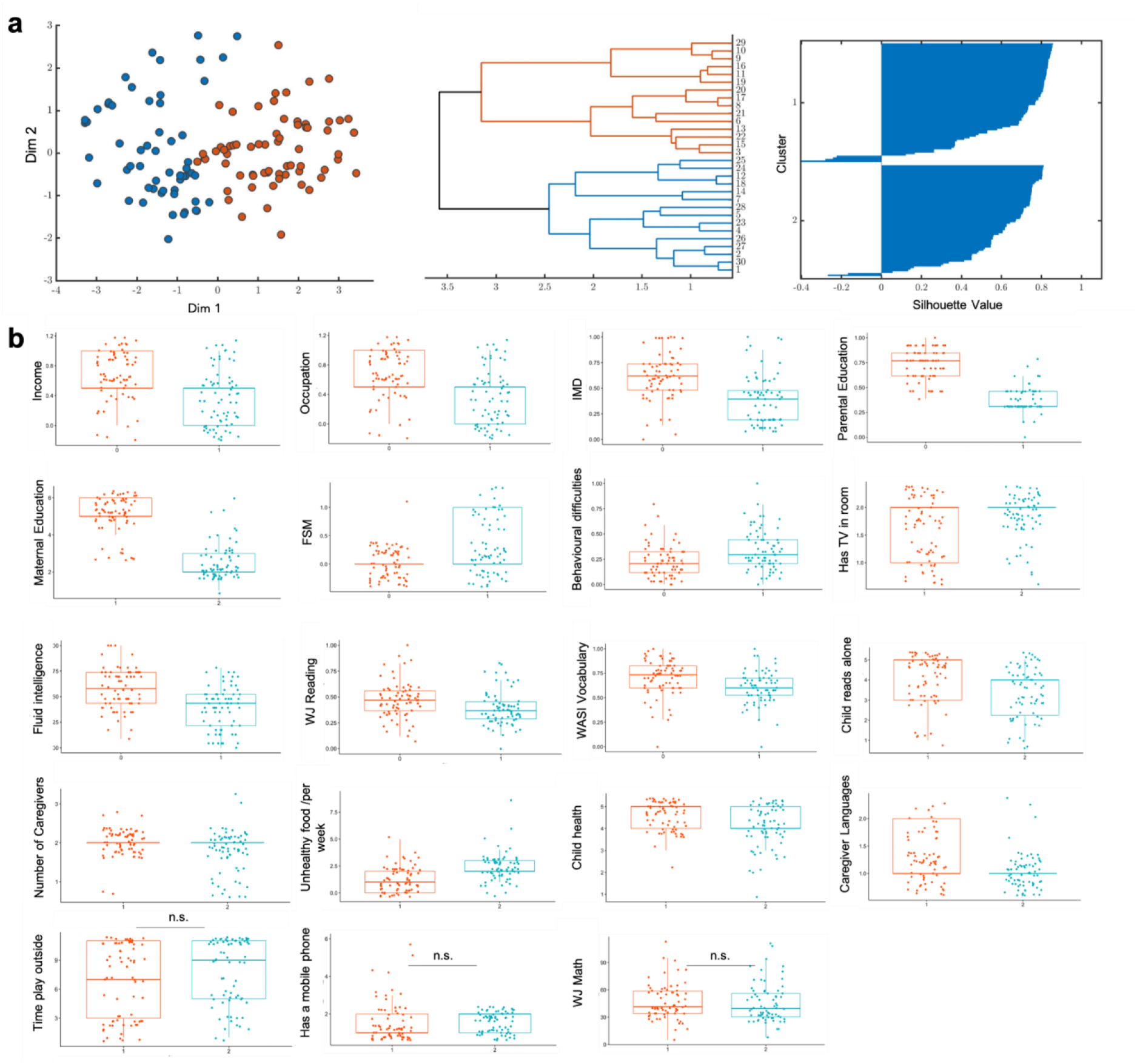
Using hierarchical clustering we defined two clearly separated SES groups and we show behavioural, cognitive and environmental differences between these SES groups. (**a**) hierarchical clustering results visualized two clusters (high SES (in red) and low SES (in blue) using scatterplot, dendrogram and Silhouette scores; (**b**) the differences between these groups/clusters are shown in many SES, environmental, behavioural and cognitive measures, only three measures were not significantly different between groups (play time outside, having or not a mobile phone and mathematical skills).

**Fig 5.**
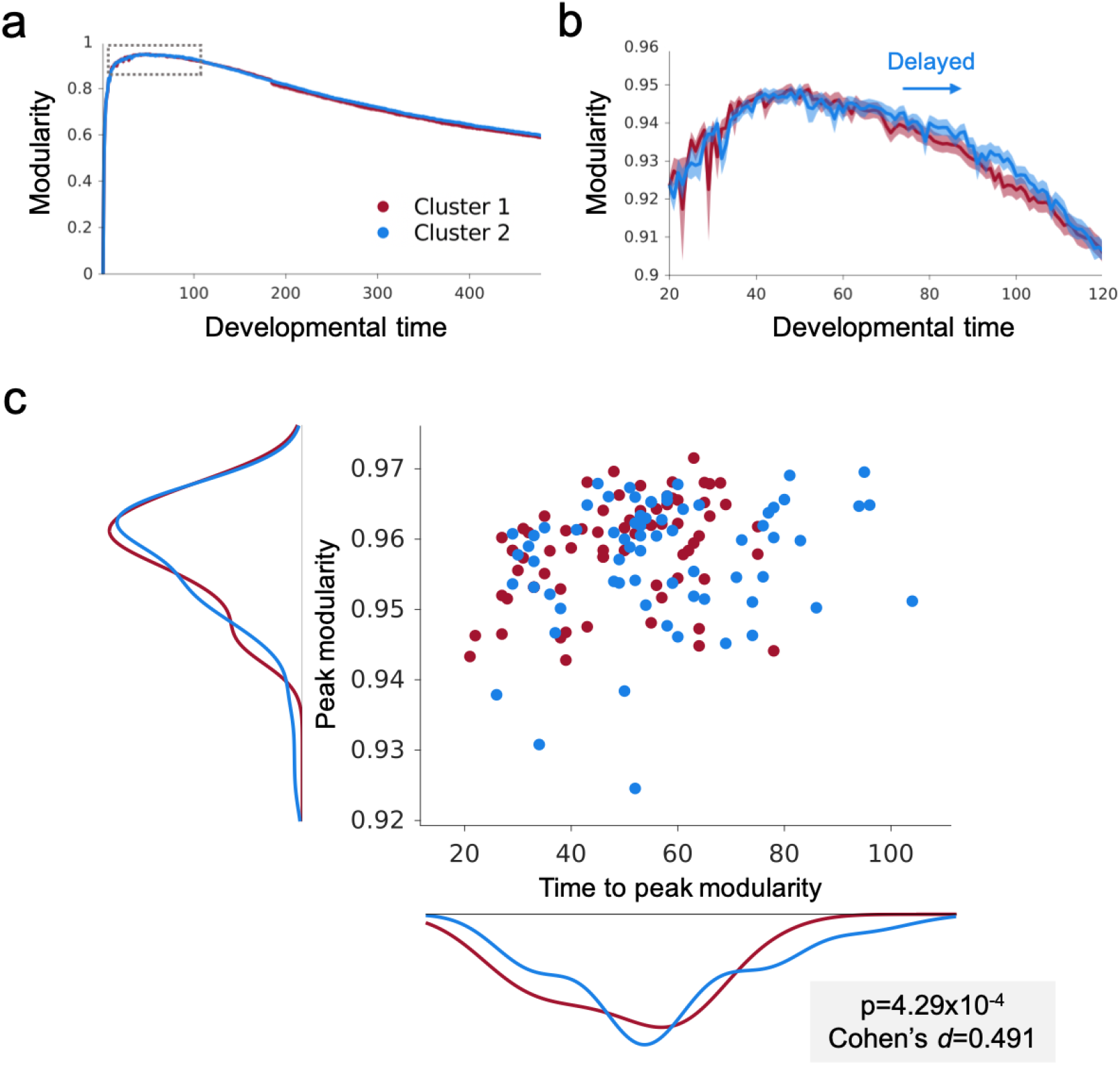
Peak modularity is delayed by shorter connection preferences. (**a-b**) The mean and standard error time courses of the two clusters are shown in terms of modularity (left) with a zoomed image of the peak modularity (right). (**c**) There are cluster differences in the time to peak modularity in Cluster 2 relative to Cluster 1. Each point reflects the peak modularity and time to peak modularity of that subject. The colour reflects their cluster allocation (Cluster 1, Red (high SES) & Cluster 2, Blue (low SES)).

### Socio-economic disadvantage and wiring parameters

Before comparing the generative models from the two groups, we first tested that they did not differ in the degree of head movement (mean framewise displacement, t_(138)_ = 0.1488, p = 0.882) or age (t_(138)_ = 1.246, p = 0.215). Another important potential confound is model energy; it is possible that the GNM better simulates the connectome formation of one cluster relative to the other, and this would confound any comparison of parameters between the clusters. However, the two groups did not differ significantly in the energy values of their best-fitting GNM (t_(138)_
 = 0.208, p = 0.836). In terms of model parameters, there were no significant differences in the cost parameter η (t_(138)_ = 0.061, p = 0.952) and the lower SES group had a lower γ parameter albeit not significantly so (t_(138)_ = 1.818, p = 0.0713).

Next, we compared the network formation trajectories of the two clusters. Children in the low SES cluster had a significantly slower time to peak modularity, relative to the higher SES cluster (*t*_(138)_ = 2.904, *P* = 4.29×10^-3^, Cohen’s d=0.491), but the height of this peak did not differ (t_(138)_ = 9,7426, *P* = 0.459). In short, the small SES-associated differences in wiring parameters significantly alter the timing network modularity emergence.

### Loosening wiring parameters increases network stochasticity

Finally, we used the mean optimal parameters taken from each cluster to simulate two sets of networks. By simulating 100 networks (50 per group) we tested systematically how the wiring probabilities (P*_ij_*) differ between the two sets of simulations. The parameters taken from Cluster 2 (low SES) produced simulated networks with more variable wiring probabilities. This was reflected both in a higher variance among wiring probabilities (Fig 6, left) and in a subtle flattening of the distribution of probability values (KS test: D = 0.012, p < 0.001) (Fig 6, right). Put simply, at each step in the simulation there are more plausible connections that might form in the brain networks of children in Cluster 2 (low SES).

**Fig 6.**
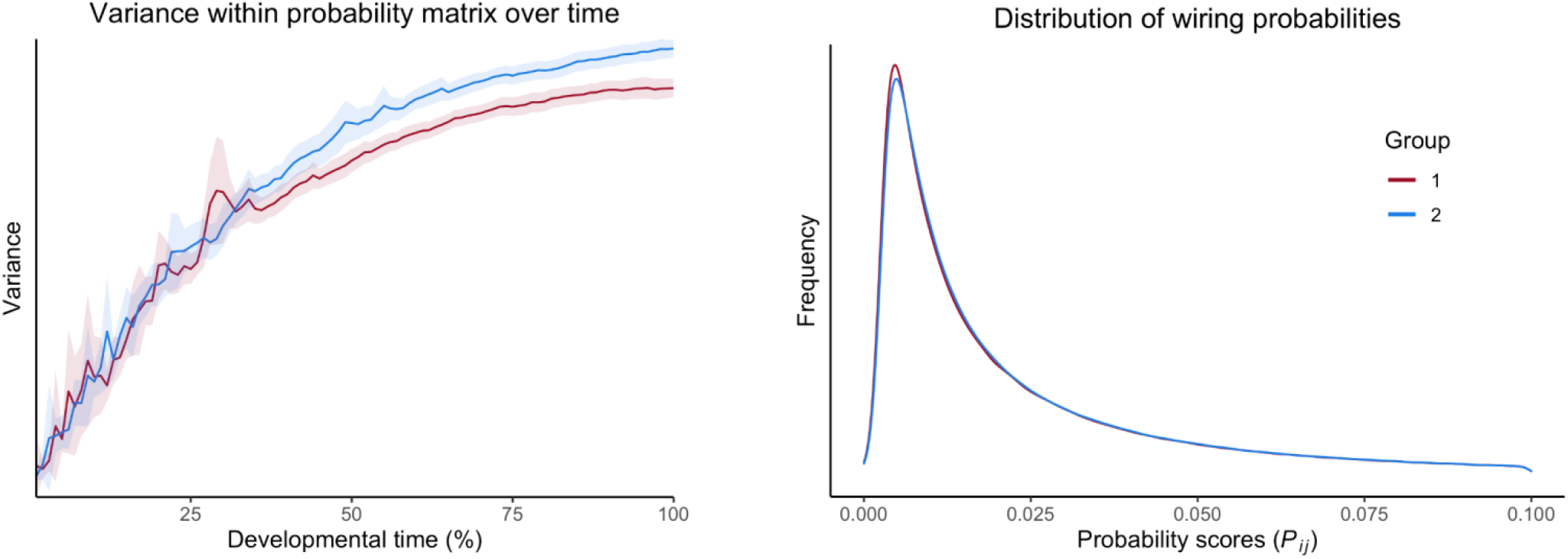
Variance over time and distribution of wiring probabilities: **Left:** Variance in values in the probability matrix (*P_ij_*) of the GNM, indicating that the wiring parameters from the second cluster produce more stochasticity in the generative process. **Right:** The distribution of the wiring probabilities within the probability matrix (*P_ij_*) of the GNM. Cluster 2 shows a flatter distribution, indicating more connections with higher wiring probabilities.

## DISCUSSION

Childhood socio-economic disadvantage is significantly associated with whole-brain network organisation [39, 57], and is associated with a host of long-term outcomes, including lower educational attainment [58], slower cognitive development [19] and poorer mental health [59]. However, the underlying mechanisms by which this disadvantage shapes neurocognitive development is not well understood. We show that, at a global scale, socio-economic disadvantage is associated with a subtle change in the mathematical rules that govern brain network formation, and that this results in a significantly altered trajectory of brain organisation, relative to that in more privileged peers. In essence, the economic constraints under which disadvantaged children’s networks form are subtly different, and in turn this impacts the emergence of an important topological characteristic, namely modularity.

### Generative networks capture the formation of observed whole-brain networks

Variability in whole-brain network formation can be simulated using a generative network which captures an iterative economic trade-off over development [11]. At the heart of this generative network is a simple wiring equation which calculates connection probabilities under different mathematical contexts. This provides a framework in which different generative principles can be formally compared and tested against observed networks. When generative networks incorporate the homophily principle – the topological value of neighbourhood similarity – within the wiring equation, they outperform all other tested generative rules (see also [13, 11]. This two-parameter equation is deceptively simple, but the iterative nature of the generative process means that network formation is modelled as a probabilistic process, with new connections altering the topological properties of the network and thus future wiring probabilities. In short, a few additional connections can guide network trajectories, because they alter neighbourhood structures and thus bring new potential connections into play [11]. This means that the simple equation produces a developmentally stochastic process.

Homophily based rules likely perform so well because they allow the network to achieve the highest computational capacity given finite biological constraints. Because nodes with shared neighbourhoods wire together, modular architectures emerge, a process observed in structural brain development [59]. It may be that homophily provides a locally-knowable heuristic – *wire with other nodes with shared neighbourhoods* – which approximates the global goal of achieving the characteristic global network organisation. This property of being ‘locally-knowable’ makes this mechanism particularly biologically plausible [60]. Indeed, it has been argued that homophily could capture Hebbian learning [61], with activity-dependent correlations providing a rough index of neighbourhood, and thus in turn driving future network connections. A stronger claim in favour of the role of homophily-based mechanisms within network formation was recently made [62], arguing that it reflects the selective reinforcement of the topological overlap within neuronal wiring and that this in turn directly contributes to the modular organisation of brain networks.

In addition to establishing the economic conditions and parameters under which brain networks grow and form trajectories of simulated development, this mathematical formalisation allows us to decompose the underlying constraints that shape network formation, and their impact on both organisational timing and network characteristics. This in turn provides rich information about individual differences in the probabilistic construction of different individual’s connectomes.

### Socio-economic disadvantage is association with an altered the trajectory of network formation

Our recruitment was biased towards oversampling children experiencing socio-economic disadvantage. Widely used measures of SES, representing standard indices of economic resources, parental education and occupation, were introduced to a data-driven clustering algorithm, producing two broad clusters that were age and gender matched. The majority of children in the second cluster lived below the poverty line, were more likely to be in receipt of free school meals, live in deprived neighbourhoods and have parents with lower educational and occupational status. The wider profile of our clusters replicate well-reported findings from the SES literature: children experiencing socio-economic disadvantage are significantly more likely to have lower cognitive performance [19, 63 – 69], educational attainment and greater behavioural difficulties [70, 71, 72], relative to their more privileged peers.

Socio-economic disadvantage was associated with small and non-significant changes in network parameters, but because these converge, they result in significantly shifted trajectories of network modularity. In particular, children experiencing socio-economic disadvantage show a slower rising peak in network modularity over the course of the simulation.

Previous studies have shown that generative network principles can be significantly associated with the genetic patterning of expression profiles [11] and polygenic risk for known neurodevelopmental conditions [73]. In the former case, a gene enrichment analysis revealed that the regional patterning of homophily values was significantly associated with cellular and biological pathways associated with synapse formation and regulation. In contrast, the regional patterning of the cost was significantly associated with cellular and biological pathways associated with metabolically costly processes, such as ribosome functioning. The implication is that individual differences in the trade-off between these two sets of biological processes provides the biophysical constraint in which complex neural networks grow, and that this subtly shifted trade-off produces differently organised neural networks across individuals.

In the current study we show that environmental factors may also shape this trade-off and its influence on network emergence. One explanation is that the brain wiring economy represents a convergent mechanism reflecting the culmination of multiple underlying biological factors that are proximal to a child’s early life environment. Based upon the extant literature on socio-economic influences on child development, we can conceive of at least three SES-related biological mechanisms that could shape wiring economy. The first of these is diet. Multiple studies have shown that dietary correlates of the wider SES construct mediate many of the established associations between socio-economic adversity and adverse health outcomes [74 – 77]. These dietary differences associated with SES – something also shown in our own sample – could impact the “cost” penalty within the wiring equation. Put simply, the relative metabolic and energetic cost of forming a connection could, at least in part, be subtly different in children according to their dietary intake. A second possible contributing factor is stress. Children experience socio-economic disadvantage also experience greater levels of stress [78 – 82], and this early stress response has been cited by many as a crucial mediating mechanistic pathway that links early life adversity to poorer long-term outcomes (e.g. [42]). One possibility is that stressful events and the resultant neuroendocrine responses alter dopaminergic neurocircuitry that supports crucial processes in the brain such as reward processing, cognitive flexibility and goal-directed behaviour [83]. These alterations could disrupt the relative trade-off between ‘costs’ and ‘values’, thereby adjusting the economic constraints within which the brain develops. The third possible contributing factor is environmental stimulation. A popular theoretical account of early childhood adversity posits that there are related but distinct dimensions of adversity which can be broadly described as ‘threat’ and ‘deprivation’ [80, 84, 85, 86]. The latter dimension corresponds most closely to socio-economic adversity, and, according to proponents of this theory, impacts early neurocognitive development through the lack of structured social, linguistic and cognitive stimulation [84]. Reduced stimulation through SES-associated differences in language environment [28, 87, 88, 89] or social context [90 – 93], could subtly adjust the relative role of the value term within the wiring equation.

In addition to delaying the peak of network modularity, the SES-associated change in wiring parameters also causes an increase in network stochasticity. In other words, at each moment in the simulation, if the parameters are weaker (as in the second cluster), then there are more plausible connections in the mix. This produces more variability in the wiring probabilities over time. On possibility is that this reflects an adaptive mechanism, rather than simply a negative consequence of low-SES. Networks that incorporate stochasticity can be more robust to certain types of perturbation [94], and in an uncertain environment heightened stochasticity may better enable the organism to adapt, albeit with poorer long-term outcomes [45].

## Conclusion

Early childhood environmental influences are associated with subtle changes in the wiring economy of the brain. As these unfold during connectome formation, the emergence of modularity is temporally shifted and network stochasticity is heightened. We propose that the economic context in which the brain develops, specifically the trade-off between the metabolic cost of a connection forming versus the topological value conferred by that connection, is shaped by multiple possible environmental factors that children experience in early life. However, it remains unclear whether this reflects a penalty incurred by networks developing in low SES environments, or a positive adaptation to environmental uncertainty.

## METHODS

### Participants

161 participants were recruited in Cambridge area, UK. The project targeted children with lower than average socio-economic status (SES) with a focussed recruitment campaign in areas with higher than average levels of community deprivation, schools with high levels of children in receipt of free school meals, and through local food banks. Participants missing more than 30% of behavioural and questionnaire data were excluded from the subsequent SES related analyses [95].

The sample included in the SES analyses comprised 145 participants, age range 6.8-12.8 (average age 9.3 years), 68 (47%) boys. Eleven percent of all children were left handed. All parents/guardians provided written informed consent, all children provided verbal assent, and the study was approved by Psychology Research Ethics Committee at the University of Cambridge (references: PRE.2015.11 and PRE.2017.102).

### Childhood outcomes

Cognitive performance was measured using the Vocabulary and Matrix Reasoning subtests from the Wechsler Abbreviated Scale of Intelligence II (WASI-II) [96]. Educational outcome measures of literacy (reading fluency) and numeracy (maths fluency) were taken from the Woodcock Johnson Tests of Achievement [97, 98]. The Strengths and Difficulties Questionnaire (SDQ) [99] with 5 categories was completed by all caregivers. This includes scales capturing peer problems, prosocial behaviour, internalising difficulties, externalising difficulties. The total difficulties score was used as an outcome measure in this study.

### Environmental measures

Primary caregivers were asked to answer questions about their child’s environment and health. Questions (Supplementary material, S1 Table) were grouped by domain (socio-economic status (SES), technology in home, early language and literacy environment, child’s health, sleep and diet).

#### a. Classic SES

For each child we had a measure that captured their parents’ highest level of primary, secondary and parental education (https://www.gov.uk/). Parental occupation was based on the specific job title, level of function and responsibility for other personnel that was later coded just on scale of (0-2, with 0 – no job, 1 – part time job, 2- full time job). We computed the average net household equivalized income, which is income after tax deductions and benefit additions, weighted by number of children and adults using Organisation for Economic Co-operation and Development (OECD) equivalence scale (Anyaegbu 2010). Average income was £28,000 (± £16,000) ranging from £3,570 to £75,000. Taking into account UK median from 2017-2018 (Office for National Statistics 2018) that was £ 31 876, 53 (36.6%) children from our sample were bellow UK poverty line (60% of the median income or less). FSM (free school meal) status was child’s eligible (in School Year 3 and above) to receipt free school meals and finally Index of Multiple Deprivation (IMD) rank measure [100]. This final measure is derived from a large-scale national administrative data based upon the family’s postcode.

#### b. Subjective SES

This self-reported measure requires caregivers to rate their current economic standing by placing them a cross on a ladder of 10 rungs, with the top representing those who were better off in the United Kingdom, and the bottom representing those the worse off. This is a frequently used measure of subjective SES called “The Macarthur Scale of Subjective Social Status” [101 – 103].

#### c. Technology in home

Each child’s access to and usage of technology, with questions including whether they have a mobile phone; the number of computers at home; availability of internet; if child has TV or computer in his room.

#### d. Early language and literacy environment

Two additional measures were added to describe child’s environment that could influence his language development: what language caregivers speak at home; and how much time child reads alone a day.

#### e. Health, sleep and diet

**Three additional questions covered** child’s overall physical health during life; the number of days a week child spends playing outside; number of unhealthy food portions per day; and, the average number of hours the child sleeps at night.

### Pre-processing of the questionnaire, behavioural and cognitive data

The data from 17 environmental, 2 cognitive, 2 educational outcomes and a behavioural measure of total difficulties from the SDQ. Fifteen participants that were excluded because they were missing more than 30% data [95]. For the small amount of remaining missing data (4% missingness overall) we used k-Nearest Neighbour imputation method based on a variation of the Gower Distance for numerical, categorical, ordered and semi-continuous variables. Gower distance is computed as the average of partial dissimilarities across individuals. Each partial dissimilarity (and thus Gower distance) ranges in [0 1] [104]. This imputation method is implemented in R (VIM package as kNN function with 5 Nearest Neighbours used) [105]. We used Spearman correlations and a Welch two sample t-test to check the quality of the kNN imputation. This compares the correlation matrices between variables of interest before and after imputation (Fig S1, Supplementary material). Correlation matrixes did not show significant differences *t*_(157.85)_ = −0.143, p =0.887.

### Hierarchical clustering of SES measures

Six SES variables (FSM, IMD, maternal education, parental education, occupation, income) were z scored and subjected to multi-dimensional scaling (using the mdscale function in Matlab). Next they were submitted to a hierarchical distance-based agglomerative clustering (using linkage in Matlab). A Silhouette analysis was conducted for each divergence point within the cluster tree, with the optimal solution being two clusters (Silhouette Coefficient = 0.62). These two clusters comprised 75 and 70 members, respectively. We subsequently used propensity matching to produce two clusters of 70 participants that were matched on age and gender.

### MRI structural scans

After being familiarised with the MRI process in the on-site mock scanner, children took part in a MRI scan whilst watching their favourite movie. All scans were obtained on the Siemens 3 T Prisma-fit system (Siemens Healthcare, Erlangen, Germany), using a 32-channel quadrature head coil. T1-weighted volume scans were acquired using a whole brain coverage 3D Magnetization Prepared Rapid Acquisition Gradient Echo (MP RAGE) sequence acquired using 1 mm isometric image resolution. Echo time was 2.98ms, and repetition time was 2250ms. Diffusion scans were acquired using echo-planar diffusion-weighted images with an isotropic set of 68 noncollinear directions, using a weighting factor of *b* = 1 000s × mm ^-2^, interleaved with 4 T2-weighted (*b* = 0) volume. 60 contiguous axial slices and isometric image resolution of 2 mm covered the whole brain. Echo time was 90ms and repetition time was 8 500ms. Children were watching their preferred movies while undergoing anatomical and diffusion scans.

### Anatomical data pre-processing pipeline

#### Anatomical data pre-processing used for diffusion data pre-processing

The T1-weighted (T1w) image was corrected for intensity non-uniformity (INU) using N4BiasFieldCorrection ([106], ANTs 2.3.1), and used as T1w-reference throughout the workflow. The T1w-reference was then skull-stripped using antsBrainExtraction.sh (ANTs 2.3.1), using OASIS as target template. Spatial normalization to the ICBM 152 Nonlinear Asymmetrical template version 2009c ([107], RRID:SCR_008796) was performed through nonlinear registration with antsRegistration (ANTs 2.3.1, RRID:SCR_004757, [108]), using brain-extracted versions of both T1w volume and template. Brain tissue segmentation of cerebrospinal fluid (CSF), white-matter (WM) and gray-matter (GM) was performed on the brain-extracted T1w using FAST (FSL 6.0.3:b862cdd5, RRID:SCR_002823, [109]).

### Diffusion data pre-processing

Any images with a b-value less than 100 s/mm^2^ were treated as a b=0 image. MP-PCA denoising as implemented in MRtrix3’s dwidenoise [110] was applied with a 5-voxel window. After MP-PCA, B1 field inhomogeneity was corrected using dwibiascorrect from MRtrix3 with the N4 algorithm [106]. After B1 bias correction, the mean intensity of the DWI series was adjusted so all the mean intensity of the b=0 images matched across each separate DWI scanning sequence.

FSL’s eddy (version 6.0.3:b862cdd5) was used for head motion correction and Eddy current correction [111]. Eddy was configured with a q-space smoothing factor of 10, a total of 5 iterations, and 1000 voxels used to estimate hyperparameters. A linear first level model and a linear second level model were used to characterize Eddy current-related spatial distortion. q-space coordinates were forcefully assigned to shells. Field offset was attempted to be separated from subject movement. Shells were aligned post-eddy. Eddy’s outlier replacement was run [112]. Data were grouped by slice, only including values from slices determined to contain at least 250 intracerebral voxels. Groups deviating by more than 4 standard deviations from the prediction had their data replaced with imputed values. Final interpolation was performed using the jac method.

Several confounding time-series were calculated based on the pre-processed DWI: framewise displacement (FD) using the implementation in Nipype (following the definitions by [113]. The head-motion estimates calculated in the correction step were also placed within the corresponding confounds file. Slicewise cross correlation was also calculated. The DWI time-series were resampled to ACPC, generating a pre-processed DWI run in ACPC space with 1mm isotropic voxels.

Many internal operations of QSIPrep use Nilearn 0.7.0 ([114], RRID: SCR_001362) and Dipy [115]. For more details of the pipeline, see the section corresponding to workflows in QSIPrep's documentation.

### DSI Studio Reconstruction

Diffusion orientation distribution functions (ODFs) were reconstructed using generalized q-sampling imaging (GQI, [116]) with a ratio of mean diffusion distance of 1.25.

### Brain network creation and measurements

After standard steps of preprocessing that are explained above, the general procedure of estimating the most probable white matter connections for each individual followed: Regions of interest (ROIs) were based on the Brainnetome parcellation of the MNI template with 105 cortical ROIs per hemisphere and 36 subcortical regions. To construct the connectivity matrix, the number of streamlines intersecting two ROIs was estimated and transformed into a density map for each pairwise combination of ROIs. A symmetric intersection was used so that streamlines starting and ending in each ROI were averaged. Self-connections were removed.

### Generative network model to grow connectomes

The generative network model (GNM) can be expressed as a simple wiring equation [12, 13, 11]. If you imagine a series of locations within the brain, at each moment in time the wiring equation calculates which two locations will become connected. It calculates this wiring probability by trading-off the cost of a connection forming, against the potential value of the connection being formed. The equation can be expressed as:

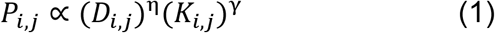

where *D_i,j_* represents the Euclidean distance between node centroids *i* and *j*, and *K_i,j_* reflects the value in forming a connection defined by the wiring rule. *P_i,j_* represents the wiring probability as a function of the product of the parameterized costs and value. The end result is a wiring probability matrix which updates over time as new connections are added.

An absolute threshold of 321 streamlines was enforced (i.e. a minimum of 321 streamlines must have connected two regions for us to consider the presence of an anatomical connection) across the sample in order to produce binary networks with mean density ≈ 2%. Across the sample, these binary networks have a mean number of connections = 603.31 (Supplementary S4 Fig.).

No seed network was used and thus the first simulated connection was generated only according to the (*D_i,j_*)^η^ term. The thirteen *K_i,j_* terms used are given in Table 2.

**Table 2.**
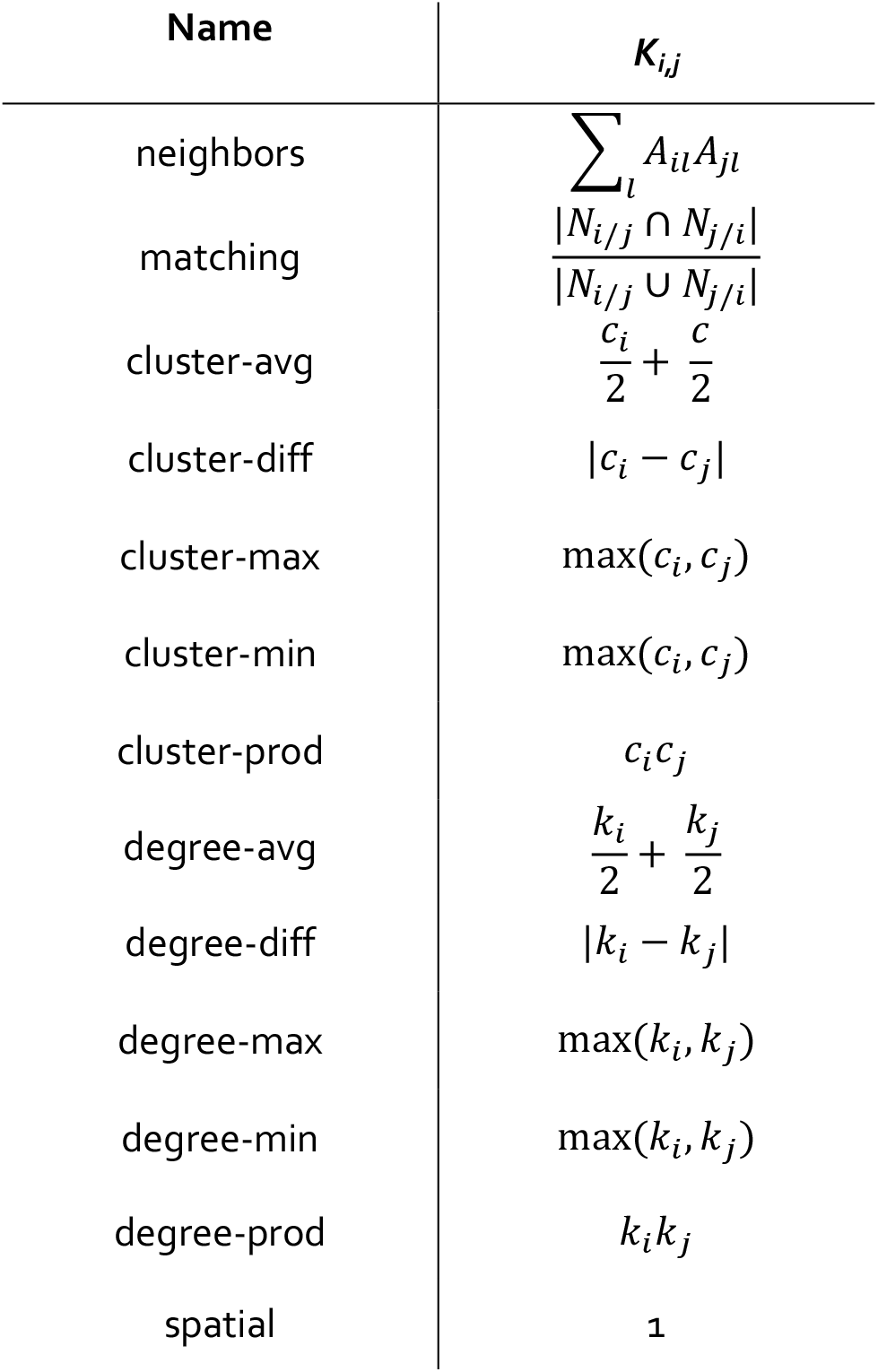
Wiring rules used within the generative network model, see Eq. 1. Node centroids were computed from brainnetome246 MNI coordinates downloaded from brainGraph v3.0.0 (https://github.com/cwatson/brainGraph). The subsequent Euclidean distance was computed with squareform(pdist(coordinates)). Across the n=145 sample, the median correlation of Euclidean distance with fiber length is r = 0.6408 for extant edges (Supplementary S5 Fig.).

### Model fitting with the Voronoi tessellation procedure

For each *η* and *γ* combination, we evaluated the model fit of the resulting network via an energy equation [12, 13]:

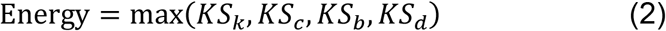

Where KS is the Kolmogorov-Smirnov statistic between the observed and simulated networks at the *η* and *γ* combination used to construct the simulation, in terms of network degree k, clustering coefficient c, betweenness centrality b and Euclidean distance d.

We optimized *η* and *γ* using a Voronoi tessellation procedure [13]. This procedure works by initially randomly sampling the parameter space and evaluating the model fits of the resulting simulated networks. On an initial batch of simulations, we simulated networks across all thirteen generative rules (as described in Table 2) with parameters taken in the range of −15 ≤ η ≤ 0 (where wiring cost is always inversely proportional to the wiring probability score) and −4 ≤ γ ≤ 4. Following an initial search of 2000 parameters, we performed a Voronoi tessellation of the space which establishes distinct two-dimensional cells of the space. We then preferentially sampled from cells with better model fits according to the energy equation, as shown in equation (2). Preference was computed with a severity of *α* = 2, which determines the extent to which cell performance led to preferential sampling in the next step. This procedure was repeated a further four times, leading to a total of 10,000 simulations being run for each network across the thirteen generative rules. On a second batch of simulations, we undertook 50,000 simulations to identify more precise optimal parameter estimates for the matching generative model, as this achieved the best model fits on the first batch of analysis. Parameters were taken in a narrower range of 4 ≤ η ≤ 0 and 0.2 ≤ γ ≤ 0.5 based on a characterization of the first batch’s parameter space. The same Voronoi hyperparameters were used as the first batch, but for 25 levels of repetitions rather than five, leading to a total of 50,000 total simulations.

### Modelling trajectories of the development

GNM offers not only the way to find a couple of brain wiring parameters that drive economy of brain network organization [9], but also dynamical changes of global organization in time that can be viewed as trajectories of brain segregation and integration. We used the max modularity as a measure of segregation and global efficiency as integration measure. Since number of modules in the brain is changing from childhood to adulthood, first increasing, reaching its maximum and then decreasing and stabilizing for a longer time, we also calculated the height of the peak, and the time to the max peak. We compared both measures in two groups (Cluster 1 and Cluster 2).

### Morphological data analysis

For ROI definition, T1-weighted images were submitted to nonlocal means denoizing in DiPy, robust brain extraction using ANTs v1.9 [117], and reconstruction in FreeSurfer v5.3 (http://surfer.nmr.mgh.harvard.edu). Regions of interests were based on the Desikan-Killiany parcellation of the MNI template [118]. The cortical parcellation was expanded by 2 mm into the subcortical white matter. The new parcellation Brainnetome246 [119] was applied afterwards using Brainnetome Atlas cortical surface files in Freesurfer. 246 ROIs (104 left side, 104 right hemisphere). From the Freesurfer analysis we used morphometric measures such as mean cortical thickness, standard deviation of cortical thickness, cortical surface area, grey volume, mean curvature, cortical folding index per each ROI.

## Acknowledgements

We would like to thank Amy Johnson and the RED Team for collecting the data included in this manuscript, they include Edwin Dalmaijer, Alex Anwyl-Irvine, Giacomo Bignardi, Tess Smith, and Stepheni Uh. Also we thank all our participants and their parents for taking part in our study. For the purpose of open access, the author has applied a Creative Commons Attribution (CC BY) licence to any Author Accepted Manuscript version arising from this submission.

