## Supplementary material for "Socio-economic disadvantage is associated with alterations in brain wiring economy": S1 Table, S2 Table, S3 Table, S1 Fig, S2 Fig, S3 Fig, S4 Fig, S5 Fig

| Variable name | Code | Values |
| --- | --- | --- |
| Age (years) | Age.Years |  |
| Gender | Child.Gender | 0, Girl  1, Boy |
| Handedness | Handedness | 1, Right  2, Left |
| Free school meal | FSM | 1, Yes  0, No |
| Subjective SES | Subjective.SES | 1 to 10 |
| Mother’s education | Education_mother_coded | 1, Primary  3, Secondary school  4, Vocational  5, Bachelor  6, Master,  7, PhD |
| Equivalized income | Equivalized.Income |  |
| The Index of Multiple Deprivation | IMD |  |
| Occupation coded | Occupation.Code | 0, No job  1, Part-time  2, Full-time |
| Caregiver speaks other languages at home | Caregiver.other.languages.at.home | 0, No  1, Yes |
| Child has a mobile phone | Child.Mobile.Phone | 0, No  1, Yes |
| No of computers at home | No.of.Computers | 1, No books  2, One book  3, Two books  4, Three books  5, Four or more |
| Child has a TV or Computer in the room | Child.Have.TV.Computer.in.the.room | 0, No  1, Yes |
| Number of books in home | Books.in.home | 1 to 4 |
| Min child read alone day (hours) | Min.child.read.alone.day | 0 to 4 |
| Total unhealthy food portions per day | Total.Unhealthy.food.perDay | 0 to 9 |
| Days a week child spends playing outside | days.a.week.child.spend.playing.outside | 1 to 7 |
| Hours child sleeps per night | hours.child.sleeps.night | 6 to 10 |

**Supplementary S1 Table.** Environmental variables codes

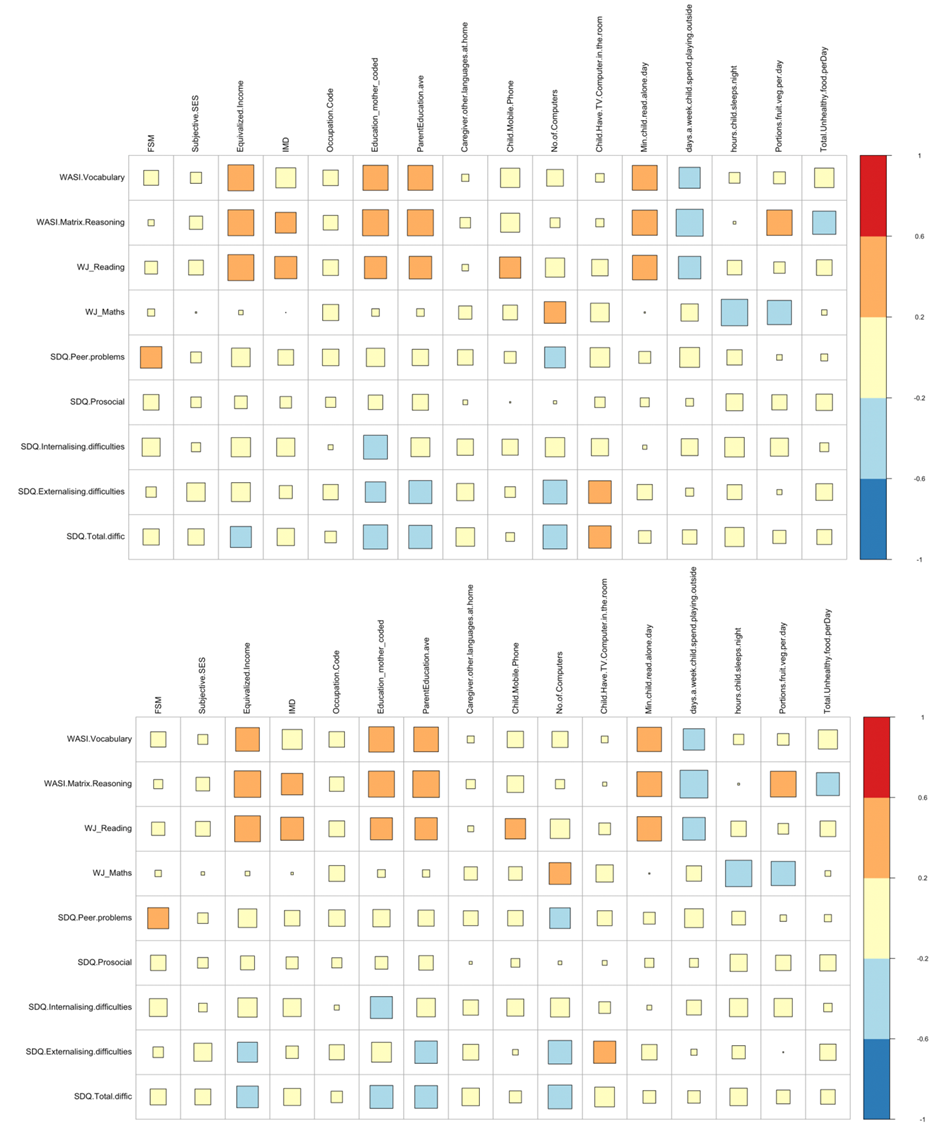

**Suplementary S1 Fig.** (**a)** Correlation between the response variables of interest in raw data with all pairwise missing values deleted. **(b)** Correlation between variables of interest after the imputation using kNN, with neighbourhood k=5.

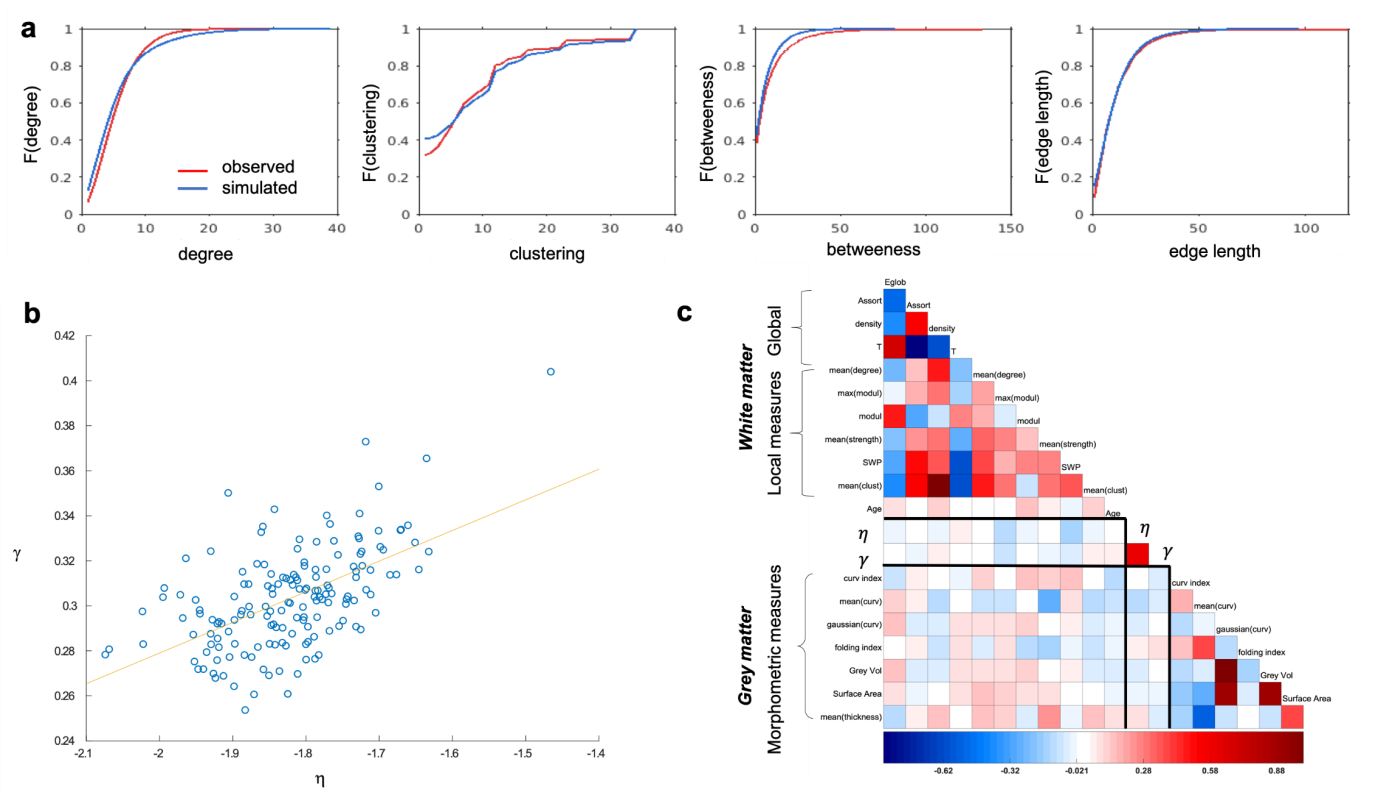
**Supplementary S2 Fig. Simulated networks grown via optimized homophily mechanisms and relationship of wiring parameters with grey and white matter measurements (on the left)** scatterplot of relationship between two wiring parameters, **(on the right)** correlation analysis of the wiring parameters with local, global and morphological measures of brain connectomes.

A

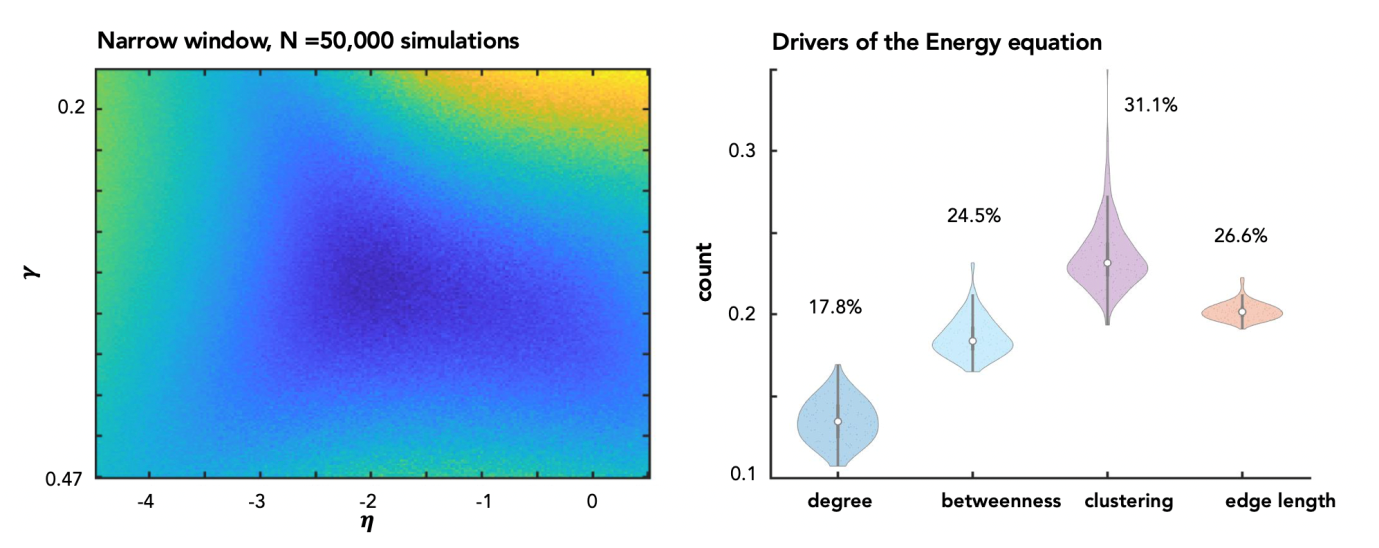

**Supplementary S3 Fig.** Parameters space with $\eta$ and $\gamma$ and the colour represents the energy of the networks they produce (blue colour represents low energy, yellow colour – higher energy values). Parameters were selected using the matching generative rule with 50,000 simulations in the limits -4 < $\eta$ < -0.5 and 0.2 < $\gamma$ < 0.5.

**Supplementary S2 Table.** Generative rules, their formulae and descriptive statistics across the broad parameter space (-15 < $\eta$ < 0, -4 < $\gamma$ < 4). Descriptive statistics of the $\eta$ and $\gamma$ required to achieve the best n=1 performing network for each subject, across the generative rules.

| **Rule** | | **Energy** | | $\boldsymbol{\eta}$ | | $\boldsymbol{\gamma}$ | |
| --- | --- | --- | --- | --- | --- | --- | --- |
| **Name** | **Class** | **Mean** | **SD** | **Mean** | **SD** | **Mean** | **SD** |
| Neighbours | Homophily | 0.0814 | 0.0101 | -1.8074 | 0.1398 | 0.2677 | 0.0244 |
| Matching | Homophily | 0.0796 | 0.0109 | -1.8474 | 0.1447 | 0.3033 | 0.0292 |
| C-Average | Clustering | 0.2165 | 0.0209 | -3.0082 | 0.1920 | -1.5350 | 1.3014 |
| C-Minimum | Clustering | 0.2449 | 0.0196 | -3.1623 | 0.2044 | -1.5403 | 1.3664 |
| C-Maximum | Clustering | 0.2154 | 0.0208 | -3.0134 | 0.2086 | -1.5806 | 1.3828 |
| C-Difference | Clustering | 0.2355 | 0.0192 | -3.2834 | 0.3082 | -0.7981 | 1.5021 |
| C-Product | Clustering | 0.2442 | 0.0186 | -3.1402 | 0.2069 | -1.4899 | 1.3298 |
| D-Average | Degree | 0.2178 | 0.0213 | -2.9842 | 0.4240 | 0.7815 | 0.2614 |
| D-Minimum | Degree | 0.2260 | 0.0185 | -3.0506 | 0.2079 | 0.1799 | 0.0682 |
| D-Maximum | Degree | 0.2216 | 0.0220 | -3.0125 | 0.4507 | 0.9475 | 0.2468 |
| D-Difference | Degree | 0.1905 | 0.0190 | -3.2413 | 0.4028 | 0.9544 | 0.6599 |
| D-Product | Degree | 0.2277 | 0.0182 | -3.0497 | 0.2236 | 0.1572 | 0.0493 |
| Spatial | Geometry | 0.2390 | 0.0196 | -3.0646 | 0.1670 | n/a | n/a |

**Supplementary S3 Table.** Descriptive statistics of high performing wiring parameters eta and gamma and the energy of the networks they produce. Parameters were selected using the matching generative rule with 50,000 simulations in the limits -4 < $\eta$ < -0.5 and 0.2 < $\gamma$ < 0.5. Parameters were averaged across a number N of high performing wiring combinations.

| **Matching: Narrow energy window** | | | |
| --- | --- | --- | --- |
| **Averaged over top N networks** | **Energy** | $\boldsymbol{\eta}$ | $\boldsymbol{\gamma}$ |
|  | **Mean (SD)** | **Mean (SD)** | **Mean (SD)** |
| 1 | 0.0736 (0.0112) | -1.7554 (0.1485) | 0.3032 (0.0248) |
| 10 | 0.0813 (0.0120) | -1.7701 (0.1040) | 0.3043 (0.0221) |
| 50 | 0.0955 (0.0145) | -1.7938 (0.0958) | 0.3029 (0.0224) |
| 100 | 0.1135 (0.0169) | -1.8278 (0.1020) | 0.2989 (0.0235) |
| 500 | 0.1262 (0.0181) | -1.8488 (0.1027) | 0.2951 (0.0233) |

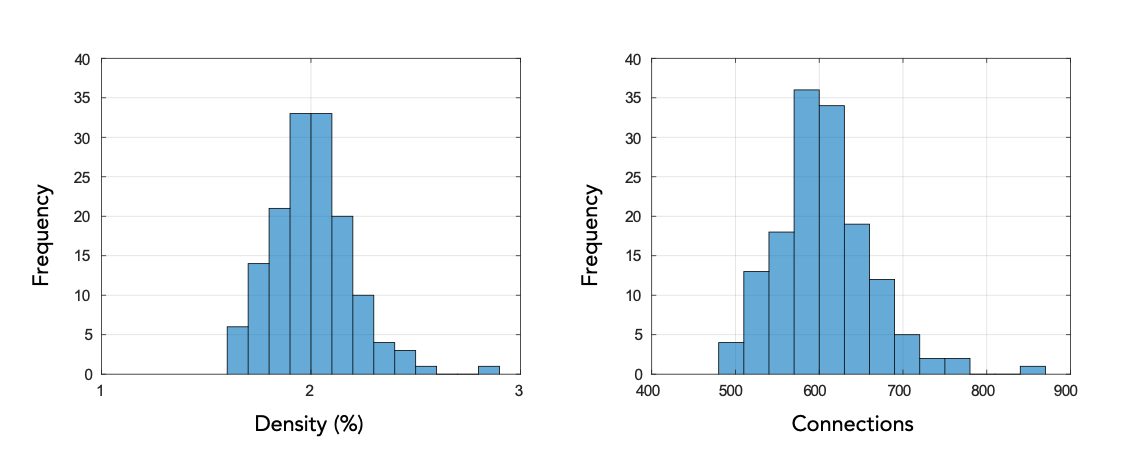

**Supplementary S4 Fig.** Histograms of the density and number of connections

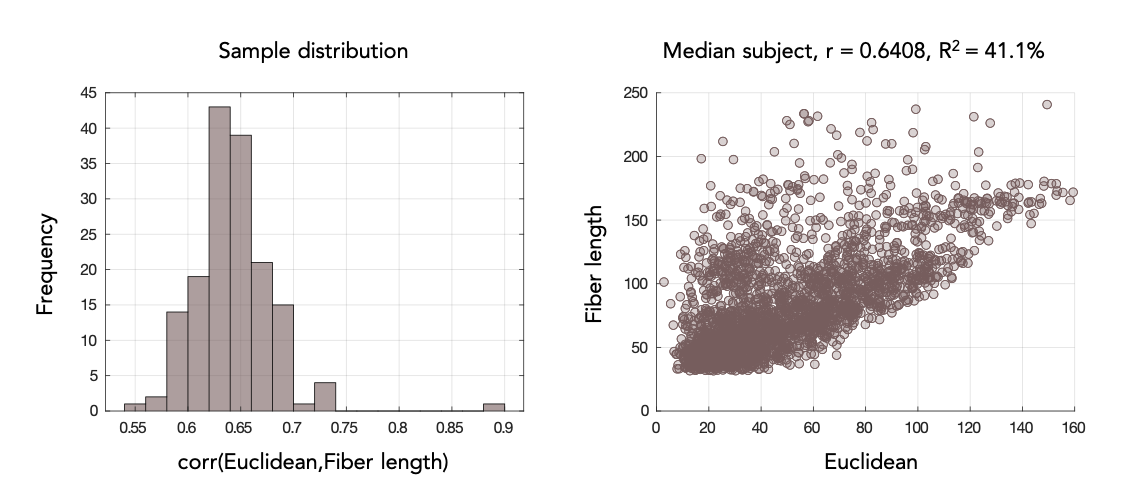

**Supplementary S5 Fig.** Correlations of Euclidean distance with fiber length. Correlation was computed using the Pearson’s rank correlation coefficient.
